# Therapeutic TNF-alpha Delivery After CRISPR Receptor Modulation in the Intervertebral Disc

**DOI:** 10.1101/2023.05.31.542947

**Authors:** Joshua D. Stover, Matthew A. R. Trone, Jacob Weston, Christian Lewis, Hunter Levis, Matthew Philippi, Michelle Zeidan, Brandon Lawrence, Robby D. Bowles

**Affiliations:** Department of Biomedical Engineering, University of Utah, Salt Lake City, UT, 84112; Department of Orthopaedics, University of Utah, Salt Lake City, UT, 84112

**Author notes:** Correspondence should be addressed to: Robby D. Bowles, Ph.D. Assistant Professor, Department of Biomedical Engineering 36 S. Wasatch Drive, SMBB 3100, Salt Lake City, UT 84112. Joshua D. Stover and Matthew A. R. Trone contributed equally to this work. **Author’s Contributions:** J.D.S., M.T., B.L., and R.D.B. all contributed to research design. J.D.S., M.T., J.W., C.L., H.L., M.P., M.Z., and R.D.B contributed to data acquisition, analysis, and interpretation. J.D.S., M.T., J.W., and R.D.B. drafted and critically reviewed the manuscript. All authors have read and reviewed the final submitted manuscript.

## Abstract

Low back pain (LBP) ranks among the leading causes of disability worldwide and generates a tremendous socioeconomic cost. Disc degeneration, a leading contributor to LBP, can be characterized by the breakdown of the extracellular matrix of the intervertebral disc (IVD), disc height loss, and inflammation. The inflammatory cytokine TNF-α has multiple pathways and has been implicated as a primary mediator of disc degeneration. We tested our ability to regulate the multiple TNF-α inflammatory signaling pathways *in vivo* utilizing CRISPR receptor modulation to slow the progression of disc degeneration in rats. Sprague-Dawley rats were treated with CRISPRi-based epigenome-editing therapeutics targeting TNFR1 and showed a decrease in behavioral pain in a disc degeneration model. Surprisingly, while treatment with the vectors alone was therapeutic, TNF-α injection itself became therapeutic after TNFR1 modulation. These results suggest direct inflammatory receptor modulation, to harness beneficial inflammatory signaling pathways, as a potent strategy for treating disc degeneration.

## INTRODUCTION

Low back pain (LBP) ranks amongst the most prevalent reasons for physician office visits, with an estimated 90% of people experiencing significant LBP at least once during their lifetime^1^. In addition, LBP is the leading cause of disability worldwide^2^, ranks third in disease burden according to disease-adjusted life years (DALYs)^3, 4^, and generates a prodigious socio-economic impact estimated at $100 billion annually in the United States alone^5^. Furthermore, disc degeneration has been implicated as the most common cause of LBP, with 40% of LBP cases attributable to the intervertebral disc (IVD)^6^. Although LBP is a malady of multifactorial origins^7–14^, and the relative importance of individual risk factors remains unknown, these risk factors initiate a cascade of molecular mechanisms leading to the pathological breakdown of the IVD.

The IVD is a cartilaginous tissue, located between vertebrae in the spinal column. The primary functions of the IVD are to confer flexibility to the spine, support mechanical loading, and provide energy dissipation. These functions are facilitated by interactions of three unique IVD tissue structures: 1) the nucleus pulposus (NP) – an inner, highly hydrated, gel-like tissue rich in aggrecan; the 2) annulus fibrosus (AF) – a tissue mainly comprised of highly organized collagen fibers circumscribing the NP; and 3) cartilaginous endplates (CEP), which border the NP and AF on the cranial and caudal surfaces, anchoring the IVDs to the adjacent vertebrae^15^. Due to the interactions between these structures, abnormal function of any one component can facilitate degeneration of the others^16, 17^. Degenerative disc disease (DDD) can be characterized by the breakdown of the extracellular matrix (ECM) of the IVD^15^, loss of disc height^18^, and inflammation^19–24^. Despite the prevalence of LBP, most treatments are palliative in nature^25^, while therapeutics that retard progression of DDD, by targeting the underlying mechanism of disc degeneration, are not available.

Pathologically elevated levels of a multitude of inflammatory cytokines have been identified in the degenerative IVD and have been implicated in the propagation of disc degeneration. TNF-α, IL-1α, IL-1β, IL-2, IL-4 IL-6, IL-10 IL-17, and interferon-γ^19, 23, 26–30^, have all been implicated in disc degeneration; however, the two most commonly studied cytokines are TNF-α and IL-1β. TNF-α signals ECM degradation, inflammation, apoptosis, and senescence^31–35^ via the NF-κβ pathway. NF-κβ is a transcription factor demonstrated to induce expression of proinflammatory and catabolic genes that drive ECM breakdown and cell survival^36, 37^. TNF-α signaling has been hypothesized to be an important driver of NF-κβ-mediated tissue breakdown, cell senescence, and apoptosis because NF-κβ inhibition has been demonstrated to ameliorate tissue breakdown in models of IVD degeneration^38, 39^. Together, these results suggest the development of therapeutics targeting the TNF-α signaling pathway as a potential treatment strategy for slowing the progression of DDD.

TNF-α modulates various cellular responses through the differential receptor profile expression of the target cells. TNF-α signaling through TNFR1 facilitates proinflammatory, catabolic, and apoptotic signaling; however, signaling through TNFR2 is anti-apoptotic and immunomodulatory in nature^35, 40^. Current treatment strategies targeting inflammatory pathways have focused on antagonism of these pathways by directly inhibiting TNF-α activity via monoclonal antibodies or other methods that broadly block cytokine activity^41^. However, these methods also eliminate TNF-α signaling through TNFR2, which may be beneficial in preventing the deleterious effects of inflammation, due to the activation of antiapoptotic and immunomodulatory signaling cascades. For these reasons, we hypothesized that therapeutic strategies capable of targeted inhibition of TNF-α/TNFR1 inflammation while simultaneously preserving the cell-protective and immunomodulatory interactions of TNF-α/TNFR2 would be more effective at slowing disc degeneration than strategies more broadly targeting TNF-α signaling. In this study, we tested this hypothesis by investigating *in vivo* CRISPRi epigenome editing of TNFR1 expression in the IVD for preventing the progression of disc degeneration in rats.

In this study, we investigated the potential of clustered regularly-interspaced short palindromic repeats (CRISPR)-based epigenome editing of inflammatory cytokine receptor profiles to slow the progression and reduce the associated pain of inflammatory cytokine-driven degenerative musculoskeletal diseases, specifically disc degeneration. CRISPR-based epigenome editing facilitates stable, highly site-specific histone modifications to modulate gene expression^42^. Briefly, CRISPR-dCas9-based epigenome editing utilizes a nuclease-deficient Cas9 (dCas9) and a customizable guide RNA (gRNA) to target specific DNA sequences in the promoter region of the target gene to regulate gene expression^43–45^. Fusion of the effector molecule Krüppel Associated Box (KRAB) to dCas9 produces highly localized H3K9 histone methylation^46–49^, which facilitates suppression of endogenous gene expression. Here, we regulate IVD cellular responses to the inflammatory environment via CRISPRi-based epigenome editing of inflammatory cytokine receptor profiles to treat disc degeneration and its associated pain in rats and show that using this approach, the response to TNA-α is altered, and even shows therapeutic effects to slow the progression of disc degeneration *in vivo*.

The objective of the present study was to test the hypothesis that CRISPR epigenome editing of inflammatory cytokine receptor profiles of IVDs can modulate cellular responses to inflammation and demonstrate CRISPRi epigenome editing of TNFR1 expression in the IVD as a potential treatment for slowing the progression of disc degeneration and treating its associated pain. This work illustrates that CRISPR epigenome editing of TNFR1 expression in the IVD slows ECM breakdown, disc height loss, inflammation in degenerative disc, and reduces pain, while switching TNF-α from deleterious to therapeutic.

## RESULTS

### Intradiscal Injection of Rat Caudal IVDs with TNFR1 Epigenome Editing CRISPRi Lentiviral (LV) Vectors Reduced TNFR1 Expression

In order to investigate the ability of CRISPRi-based epigenome editing therapeutics to regulate TNFR1 expression in the IVD, we first demonstrated the successful *in vivo* delivery and expression of LV vectors in the IVD. Toward this end, we measured the biodistribution of LV vector expression following rat caudal IVD injections via *ex vivo* bioluminescent imaging. Luciferase-expressing LV vectors were delivered to rat caudal IVDs via fluoroscope-guided intradiscal injection (**Figure 1A**). One week after injections, *ex vivo* bioluminescent imaging^50^ was conducted to quantify transgene (luciferase) expression in IVDs, surrounding tissue, and vital organs. IVDs injected with luciferase-expressing LV vectors exhibited robust bioluminescent signal confined to injected IVDs (**Figure 1B, C**). In addition, luciferase expression was not detected in control IVDs, kidneys, liver, spleen, heart, or lungs (**Figure 1C**). Together, these results demonstrated our ability to sequester LV vector delivery and expression within the target IVD.

**Figure 1.**
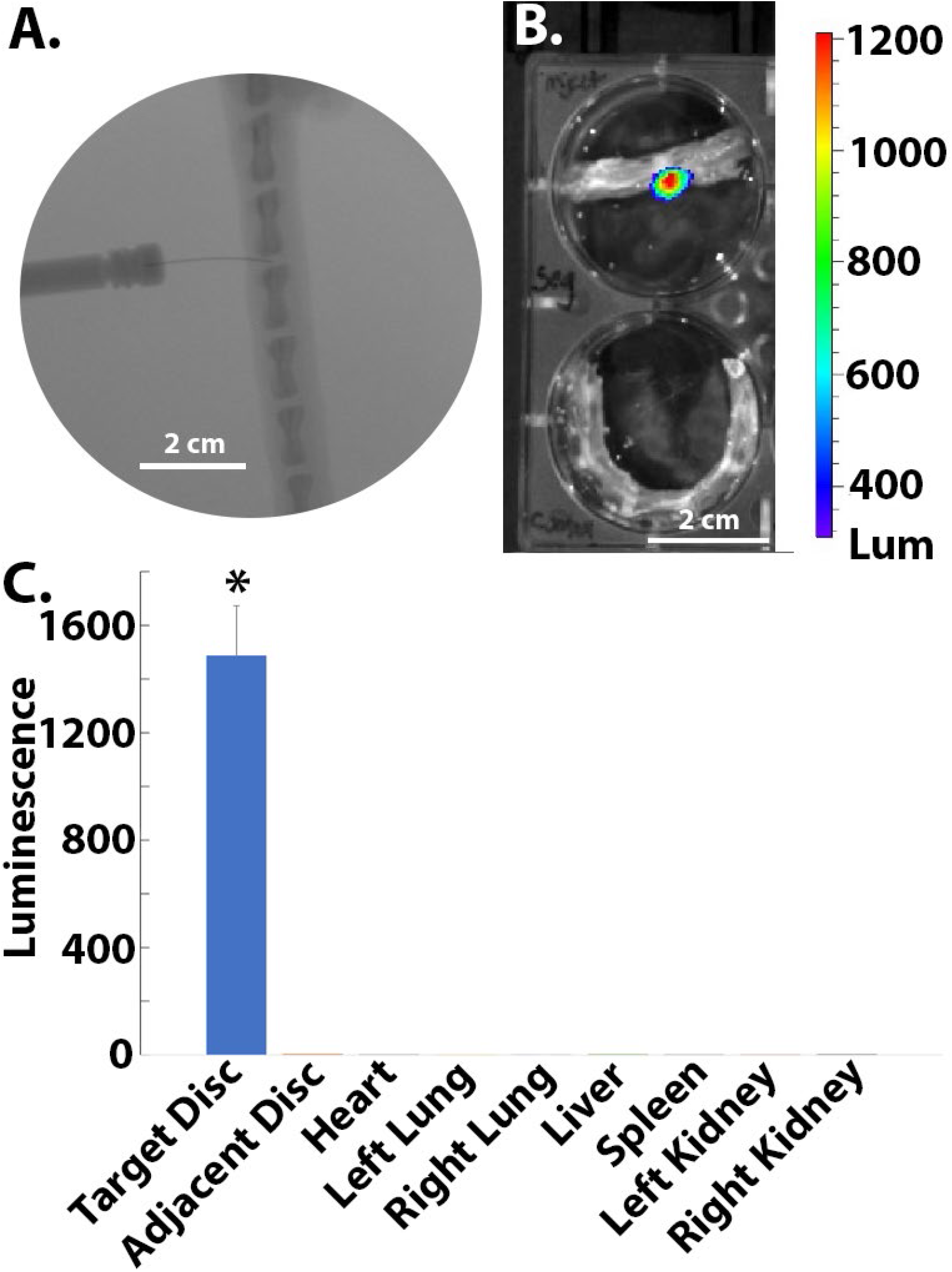
Biodistribution of lentiviral vector expression is sequestered within target intervertebral discs (IVDs) following intradiscal injection of rat caudal IVDs. **A).** Rat caudal IVDs were injected via fluoroscope-guided intradiscal injection with luciferase expressing lentiviral (LV) vectors. Lentivirus expression was measured via *ex vivo* IVIS bioluminescence imaging 1 week after injection. **B).** *Ex Vivo* bioluminescence images of luciferase-expressing LV vector injected (top) and control (bottom) rat tail segments demonstrated LV expression sequestered within target disc. Red: high LV expression; Blue: low LV expression; Clear: no LV expression. **C).** Quantification of bioluminescent signal intensity demonstrated biodistribution of LV vector expression is localized to target (injected) IVDs.; n= 5 animals; * denotes p<0.05 compared to control disc.

Once we demonstrated localized expression of LV vectors to the IVD, we tested the ability of our CRISPRi epigenome editing-based therapeutics to regulate TNFR1 protein expression *in vivo* without disrupting healthy IVD structure. Previously, CRISPRi based epigenome editing vectors targeting the gene promoter region of TNFR1 were designed, built, and their ability to regulate TNFR1 gene expression *in vitro* was demonstrated^51, 52^. In this study, rat caudal IVDs received intradiscal injection of saline (PBS), non-target, or TNFR1-targeting CRISPRi epigenome-editing lentiviral vectors (**Figure 2A**). Three weeks following injections, animals were sacrificed and hematoxylin-eosin (H&E), Alcian blue, and TNFR1 immunohistochemistry (IHC) staining of rat caudal IVD histology sections were conducted and analyzed (**Figure 2A**).

**Figure 2.**
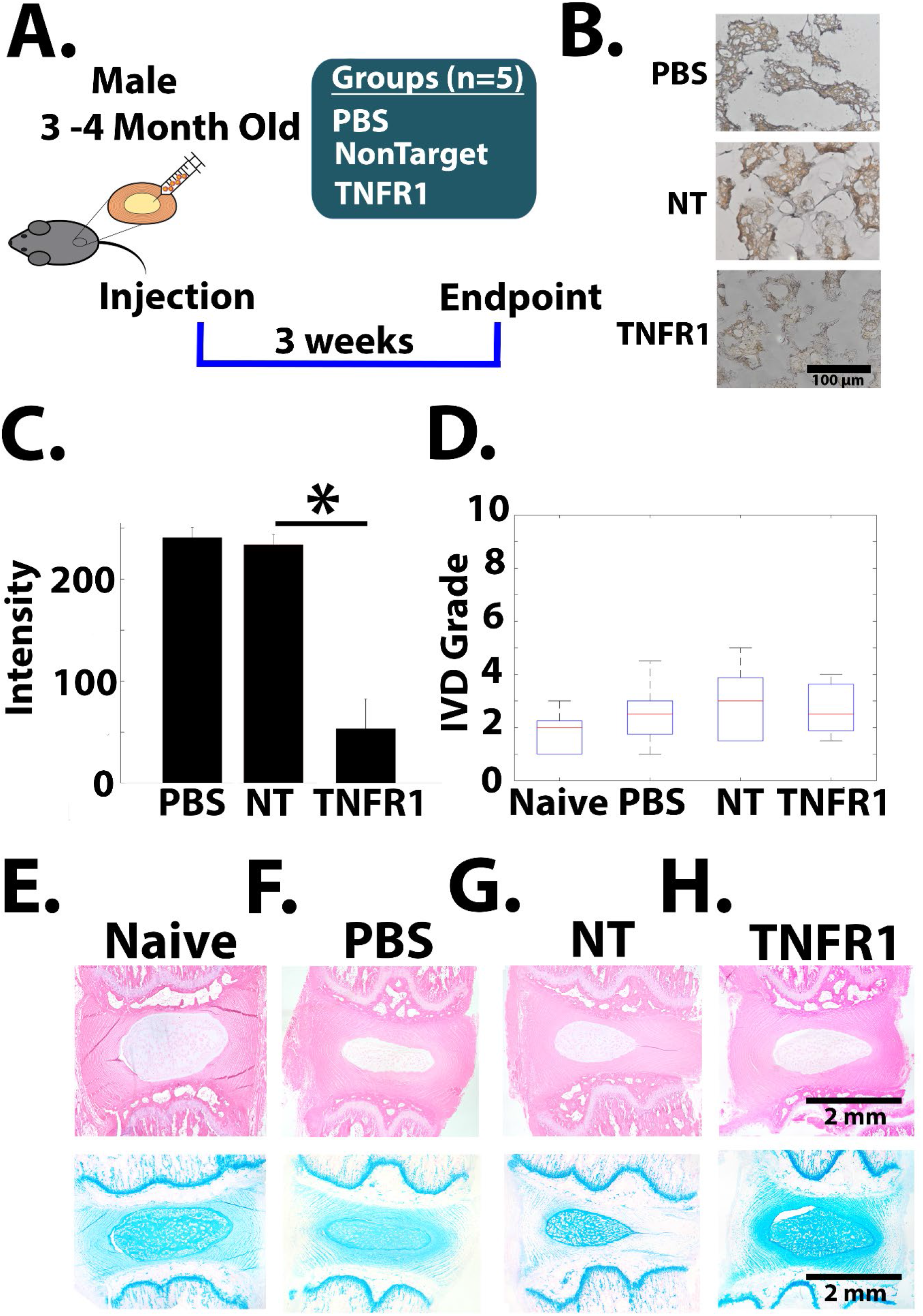
*In vivo* CRISPR based epigenome editing of TNFR1 in rat caudal IVDs robustly downregulates TNFR1 expression while maintaining healthy IVD morphology. **A).** A schematic of these experiments: 3-month-old Sprague Dawley rats received intradiscal injection of PBS, nontarget, or TNFR1 epigenome editing LV vectors (n=5 animals/groups). 3 weeks following injection, animals were sacrificed, and IVD histology, degeneration grading, and IHC for TNFR1 were performed. Naïve discs obtained from adjacent segment of PBS-injected animals (n=5). **B).** Immunohistochemistry (IHC) staining of TNFR1 expression (brown) in rat NP cells in representative IVD images from animals in the PBS (top)), non-target epigenome editing (**NT**) (middle), and TNFR1 epigenome editing (**TNFR1**) (bottom) CRISPR lentiviral vector injection groups. **C).** Quantification of TNFR1 IHC staining intensity from IHC images demonstrated significantly downregulated TNFR1 expression in IVDs from animals in the TNFR1 epigenome edited discs when compared to IVDs from animals in the PBS (PBS), and non-target (TNFR1) epigenome editing groups; * denotes p< 0.05 compared to PBS treatment group. **D).** Degeneration scoring of histology images from animals in the naïve, PBS, nontarget epigenome editing, and TNFR1 epigenome editing groups demonstrated no significant differences between treatment groups; Images scored by blinded observers**. E-H).** Images of hematoxylin-eosin (H&E) (top) and Alcian blue (bottom) staining of histological sections of IVDs from animals in the naïve, PBS injection, nontarget epigenome editing and TNFR1 epigenome-editing LV vector injection groups. Histology demonstrated preservation of tissue architecture, cell morphology, and proteoglycan content in the nucleus pulposus (NP), annulus fibrosus (AF), and cartilage end plate (EP) of TNFR1 epigenome-edited (TNFR1) IVDs;

TNFR1 expression was observed in IVDs from the PBS, non-target, and TNFR1-targeting epigenome editing lentiviral vector injection groups (**Figure 2B, brown**). TNFR1 staining intensity in IVDs from the TNFR1 epigenome-edited group (53.2±29.1) was significantly reduced when compared to staining intensity in IVDs from the PBS (240.6±5.21), and non-target epigenome editing groups (234±10.1) (**Figure 2C**), indicating reduced TNFR1 expression. In addition, IVD degeneration scores of TNFR1 epigenome-edited (2.7±1.0) IVDs were not significantly different from degeneration scores of IVDs from the naïve (1.8±0.8), PBS (2.5±1.3) and non-target epigenome-edited (2.9±1.5) groups (**Figure 2D**), indicating downregulation of TNFR1 had no deleterious effects on IVD structure in the healthy IVD. Moreover, H&E and Alcian blue stained histology sections of TNFR1 epigenome-edited IVDs exhibited typical IVD tissue architecture, disc height, and proteoglycan content (**Figure 2H**) indicative of healthy IVDs. Similarly, IVDs in the naïve (**Figure 2E**), PBS (**Figure 2F**), and non-target epigenome edited (**Figure 2G**) treatment groups maintained healthy IVD structure, following injection. Together, these results demonstrate epigenome editing of the TNFR1 gene promoter of rat caudal IVDs *in vivo* regulates TNFR1 expression while maintaining healthy IVD structure and morphology. Moreover, no intraoperative complications, signs of discomfort, or adverse health outcomes were observed in rats from all treatment groups, indicating delivery and expression of CRISPRi epigenome editing LV vectors to caudal IVDs were well tolerated.

### *In Vivo* CRISPRi Based Epigenome Editing of TNFR1 Gene Promoter in Rat Caudal IVDs Maintained Long-Term TNFR1 Downregulation and Slowed Long-Term Small Needle Puncture-Induced Disc Degeneration

Once we demonstrated our ability to deliver TNFR1 epigenome editing LV vectors to the IVD and to regulate TNFR1 expression *in vivo,* we investigated the ability of TNFR1 epigenome editing vectors to maintain long-term downregulation of TNFR1 expression *in vivo*. Rat caudal IVDs were injected, via 31-G needle (size previously demonstrated in literature to not induce degeneration for up to 4 weeks post-puncture^53^) with PBS, non-targeting CRISPR epigenome editing LV vectors, or TNFR1-targeting CRISPR epigenome-editing LV vectors. Three months following injection, TNFR1 expression and disc degeneration were evaluated via analysis of histological sections of IVDs from animals in all treatment groups (**Figure 3A**). TNFR1 staining intensity for animals in the PBS (188.6±15.6; **Figure 3B, C**) and non-target (203±13.3; **Figure 3B, C**) epigenome editing LV vector injection groups were significantly elevated over naïve controls (144.2±15.5; **Figure 3B, C**); however, TNFR1 staining intensity (110±16.4; **Figure 3B, C**) was significantly (p<0.05) decreased in animals from the TNFR1 epigenome-edited treatment group when compared to all other treatment groups (**Figure3C**). These results demonstrate that intradiscal injection of TNFR1 epigenome-editing vectors provided long-term downregulation of TNFR1 expression in the IVD.

**Figure 3.**
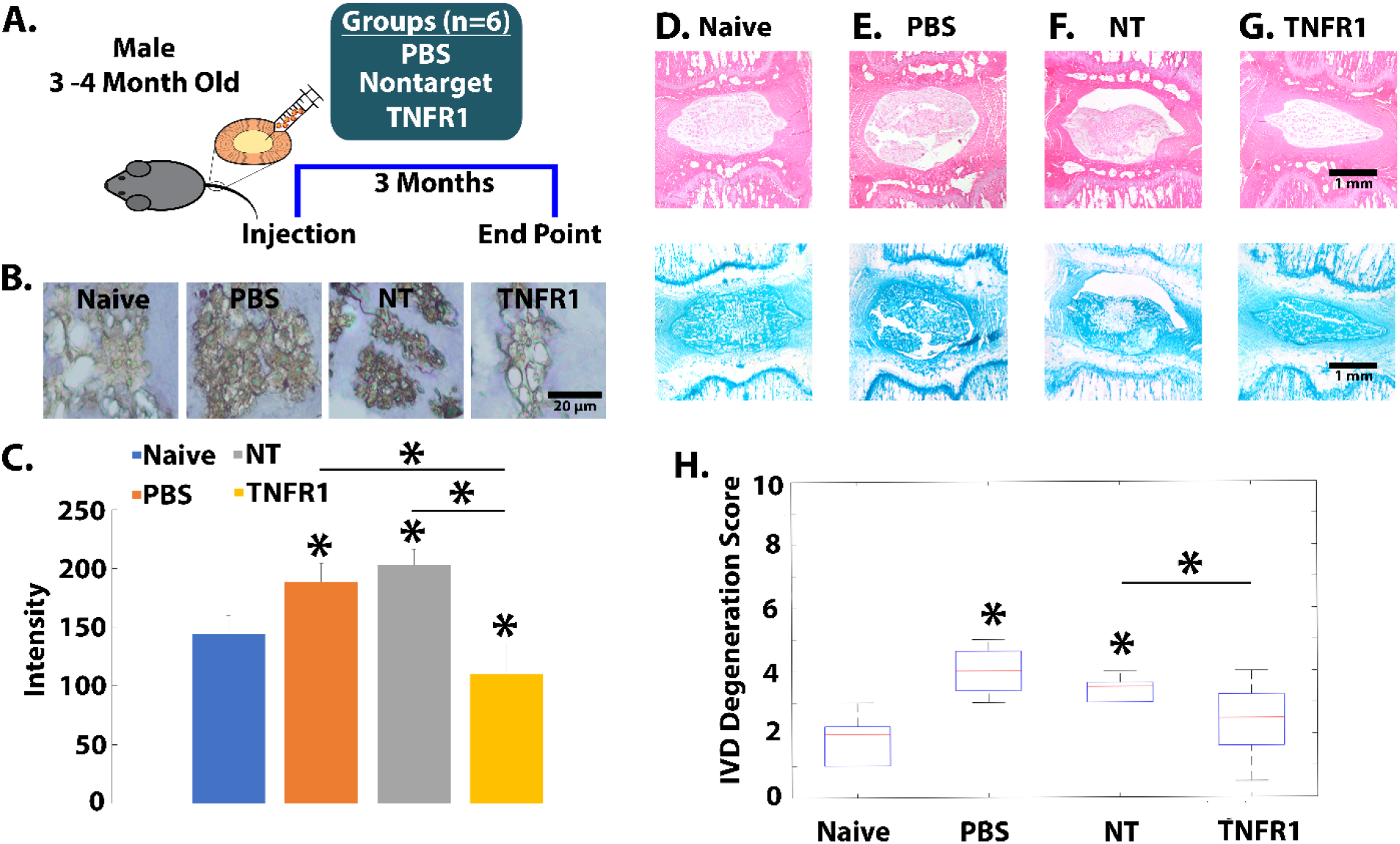
*In vivo* CRISPR-based epigenome editing of TNFR1 expression in rat caudal IVDs slows small needle puncture-induced disc degeneration and maintains TNFR1 – downregulation for 3 months post-injection. **A).** A schematic of the study design: 3-month-old, male, Sprague-Dawley rats received no injection, or injection of PBS, nontarget (NT) epigenome editing, or TNFR1 (TNFR1) epigenome editing LV vectors. 3 months post-injection, animals were sacrificed, and histology, IVD degeneration scoring, and IHC for TNFR1 were performed on IVDs from animals in each treatment group (n= 6 animals/group). **B).** Immunohistochemistry (IHC) staining of TNFR1 expression (brown) in NP cells in representative IVD images acquired from animals in the naïve (left), PBS injected (middle left), non-target epigenome editing LV vector injected (middle right), and TNFR1 epigenome editing LV vector injected treatment groups (right). **C).** Quantification of TNFR1 staining intensity revealed significantly upregulated TNFR1 expression in the PBS, and non-target epigenome edited groups; however, TNFR1 expression was significantly downregulated in the TNFR1 epigenome edited group; * denotes p< 0.05 compared to naïve (no injection) group. Images of H&E (top) and Alcian blue (bottom) of histological sections of IVDs from animals in the **D).** naïve, **E)**. PBS injection, **F).** non-target, and **G)**. TNFR1 epigenome editing LV vector injection treatment groups. **D-G).** Histology demonstrated preservation of tissue architecture, cell morphology, and proteoglycan content in the AF, NP, and endplates of naïve **(D.)** and TNFR1 epigenome edited IVDs **(G.)**; however, AF, NP, and endplates in the PBS (**E.**), and non-target epigenome editing group (**F.**) exhibited changes to tissue architecture consistent with disc degeneration **H).** Degeneration scoring (0-10) of histological images of IVDs from animals in the naïve, PBS injected, non-target epigenome editing LV vector injected and TNFR1 epigenome editing LV vector injected treatment groups revealed significantly elevated degeneration scores in the PBS and NT treatment groups; however, the degeneration scores of TNFR1 epigenome edited IVDs were not significantly different from naïve (healthy) degeneration scores * denotes p<0.05 compared to naïve (no injection) group.

IVDs from animals in the naïve (**Figure 3D**) and TNFR1 epigenome editing treatment groups exhibited healthy IVD morphology (**Figure 3G**); however, IVDs from animals in the PBS (**Figure 3E**) and non-targeting epigenome editing groups (**Figure 3F**) exhibited subtle but significant changes to IVD tissue morphology consistent with IVD degeneration. In addition, TNFR1 epigenome-edited IVDs maintained degeneration scores not significantly different from naïve IVD degeneration scores (**Figure 3H**); however, degeneration scores from IVDs in the PBS and non-target epigenome-editing treatment group were significantly elevated over naïve controls (**Figure 3H**). Together, these results demonstrate CRISPRi based epigenome editing of the TNFR1 gene promoter produces robust, long-term downregulation of TNFR1 expression and slows the progression of chronic disc degeneration in rats. Moreover, these results indicated small needle puncture of IVDs, previously shown not to induce acute disc degeneration, produce chronic disc degeneration 3 months following puncture.

### *In Vivo* CRISPRi-Based Epigenome Editing of TNFR1 Expression in Rat Caudal IVDs Significantly Decreased Disc Height Loss, Slowed Disc Degeneration, and Reduced Inflammation in an Injury Model of Disc Degeneration

To further investigate the therapeutic potential of CRISPR-based TNFR1 epigenome-editing vectors, we tested their ability to prevent disc degeneration in *in vivo* rodent models of disc degeneration. As above, rat caudal IVDs were injected with either PBS, non-targeting, or TNFR1-targeting epigenome-editing LV vectors (**Figure 4A**). Three weeks after injection, disc degeneration was induced via annular puncture (AP). Four weeks following AP, disc height loss, disc degeneration, and intradiscal inflammatory cytokine levels were measured (**Figure 4A**).

**Figure 4.**
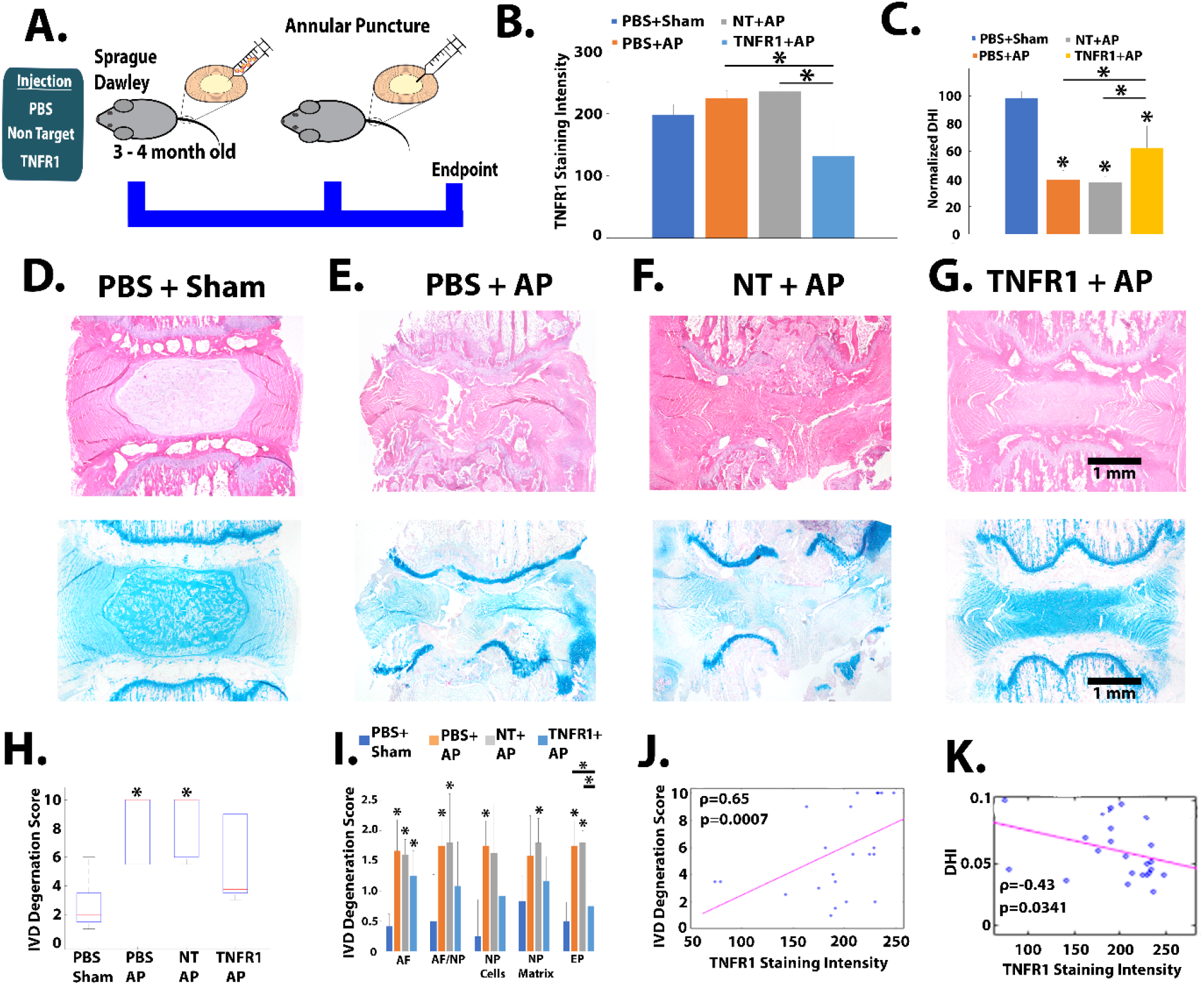
*In vivo* CRISPR-based epigenome editing of TNFR1 expression in rat caudal IVDs slows progression of disc degeneration and reduces disc height loss in annular puncture model of disc degeneration. **A).** Schematic of study design: 3-month-old male, Sprague-Dawley rats received caudal intradiscal injection of PBS (n=12 animals), non-target (NT) (n=6 animals) or TNFR1 epigenome editing LV vectors (TNFR1). Three weeks post-injection PBS-injected animals received either sham (n=6) or annular puncture (AP) (n=6), while all NT and TNFR1 received annular puncture. Four weeks following AP, disc height images were obtained, animals were sacrificed, and TNFR1 IHC, histology, and degeneration scoring of IVDs were conducted. **B).** Quantification of TNFR1 IHC staining intensity of NP cells in IVDs from animals in the PBS injection and sham, (PBS+sham), PBS injection and AP (PBS+AP), NT injection and AP (NT+AP), and TNFR1 injection and AP (TNFR1+AP) treatment groups; * denotes p<0.05 compared to sham**. C).** Disc height index (DHI) calculated from before/after AP fluoroscopic images of rat caudal IVDs of animals in the PBS+sham, PBS+AP, NT+AP and TNFR1+AP treatment groups. * denotes p<0.05 compared to PBS+sham group. **D-G).** Images of H&E (top) and Alcian blue (bottom) stained histology sections of rat caudal IVDs from animals in the **D.**) PBS+sham, **E.**) PBS+AP, **F.**) NT+AP, and **G).** TNFR1+AP treatment groups. IVD degeneration scores of **H).** total IVDs and **I)** by disc region; annulus fibrosus (AF), nucleus pulposus (NP), AF and NP border (AF/NP), NP cellularity (NP Cell), NP matrix, and endplate (EP); *denotes p<0.05 compared to PBS+sham group. Spearman correlation modeling of **J).** IVD degeneration score and **K).** DHI as a function of TNFR1 staining intensity.

First, we confirmed the ability of TNFR1-targeting CRISPRi epigenome editing LV vectors to modulate TNFR1 expression, following AP, in rat caudal IVDs. TNFR1 staining intensity of TNFR1 epigenome-edited IVDs in the AP treatment group (TNFR1+AP) (**Figure 4B**) was significantly (p <0.05) decreased when compared to TNFR1 staining intensity in the sham (PBS + sham) and non-target epigenome-edited AP (NT+AP) treatment groups (**Figure 4B**). Together, these results demonstrate treatment with CRISPRi-based epigenome-editing therapeutics targeting the TNFR1 gene promoter maintained downregulation of TNFR1 expression throughout the experiment in the aftermath of AP.

Next, we investigated the ability of TNFR1 epigenome-editing therapeutics to prevent disc height loss during degeneration. Disc height index (DHI) was calculated from radiographic images of caudal IVDs obtained at time of AP and at study endpoint (4 weeks post-AP). IVDs from rats in the PBS+AP and NT+AP treatment groups exhibited significantly reduced (p<0.05) DHI (39.5±6% and 37.6±4%, respectively, **Figure 4C**) when compared to the DHI from animals in the PBS+sham group (98.4±11%, **Figure 4C**). DHI from animals in the TNFR1+AP group (62.4±13%) exhibited significant disc height recovery when compared to the DHI of animals from the NT+AP treatment group (37.6±4%, **Figure 4C**); however, DHI remained significantly decreased (p<0.05) compared to baseline.

In addition to disc height loss, IVD morphology was evaluated from H&E and Alcian blue stained IVD histology sections. IVDs from animals in the PBS+sham _group_ (**Figure 4D**) exhibited healthy IVD tissue architecture characterized by a well-defined NP and intact NP/AF border, well-defined lamellae in the AF, normal disc height, intact end plates, healthy NP cellularity, and robust proteoglycan (PG) expression in both NP and AF tissue (**Figure 4D**). IVDs from animals in the PBS+AP (**Figure 4E**) and NT+AP (**Figure 4F**) groups exhibited hallmark characteristics of degeneration including end plate fracture, disc height loss, the loss of NP cellularity and organization, disruption of NP/AF border, loss of lamellae structure in the AF, and loss of PG expression in NP and AF tissue. IVDs in the TNFR1+AP treatment group (**Figure 4G**) exhibited recovered disc height, maintained end plate integrity, improved AF and NP organization, and restored PG expression in the NP and AF tissue (**Figure 4G**).

Furthermore, disc degeneration scores were generated by analysis of histology images by blinded observers. IVDs from animals in PBS+sham group exhibited healthy IVD morphology and received low IVD degeneration scores (2.7±1.7, **Figure 4H**). Degeneration scores of IVDs from animals in the PBS+AP and NT+AP treatment groups were (8.5±2.3 and 8.6±2.2 respectively, **Figure 4H**) significantly (p<0.05) elevated over baseline control (PBS+sham), while IVDs from animals in the TNFR1+AP treatment were not (5.3±2.9, **Figure 4I**).

Examination of disc degeneration by tissue region revealed AF degeneration scores were significantly elevated over sham in all treatment groups (**Figure 4I**). Disc degeneration scores for the NP cellularity, NP matrix, and endplate were significantly elevated in IVDs from animals in the PBS+AP or NT+AP treatment groups or both (**Figure 4I**); however, these degeneration scores were returned to baseline levels in the TNFR1+AP treatment group (**Figure 4I**). Moreover, Spearman correlation modeling of IVD degeneration scores (**Figure 4J**) and DHI (**Figure 4K**) as a function of TNFR1 expression demonstrated both IVD degeneration scores and DHI are correlated with TNFR1 expression.

In addition to the breakdown of disc structure and the loss of disc height, degenerative discs exhibit pathologically elevated levels of inflammatory cytokines^54^. In particular, the inflammatory cytokines TNF-α and IL-1β have been implicated as primary drivers of disc degeneration^31, 55^ while IL-6 has been implicated in models of discogenic pain^23, 56–58^. Based on these findings, we tested the ability of TNFR1 epigenome editing of rat IVDs to modulate the inflammatory response during disc degeneration.

Immunohistochemistry staining of TNF-α in histology sections of IVDs from animals in the PBS+AP and NT+AP treatment groups exhibited increased TNF-α positive staining (**Figure 5A**) when compared to the PBS+sham and TNFR1+AP groups (**Figure 5A**). In addition, the percentages of TNF-α immunopositive cells in IVDs from animals in the PBS+AP and NT+AP group (**Figure 5B**) were significantly elevated (p<0.05) over the percentages of TNF-α immunopositive cells in the PBS+AP group (**Figure 5B**); however, the percentage of TNF-α immunopositive cells were significantly reduced in the TNFR1+AP treatment group (**Figure 5B**).

**Figure 5.**
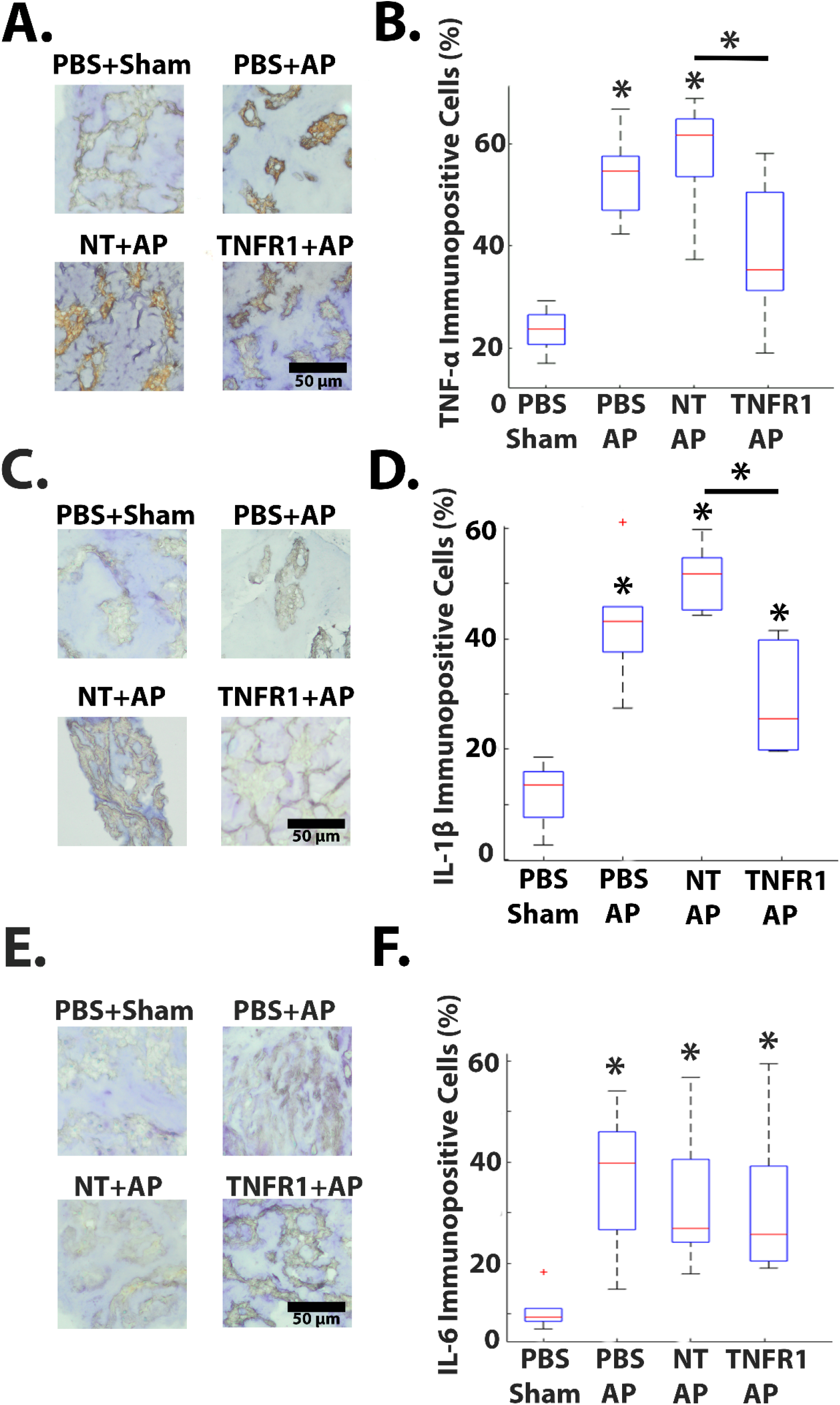
*In vivo* CRISPR epigenome editing of TNFR1 expression in rat caudal IVDs reduced expression of TNF-α, and IL-1β in annular puncture model of disc degeneration. **A).** Images of immunohistochemistry (IHC) staining for TNF-α expression (brown) in rat NP cells from IVDs in the PBS injection and sham (**PBS+sham**) (top left), PBS injection and annular puncture (AP) (**PBS+AP**) (top right), non-target epigenome editing LV vector injection and AP (**NT+AP**) (bottom left), and TNFR1 epigenome editing LV vector injection and AP (**TNFR1+AP**) (bottom right) treatment groups. **B).** Percentage of TNF-α immunopositive NP cells; *denotes p<0.05 compared to PBS+sham. **C).** Images of IHC staining for IL-1β expression (brown) in rat NP cells in the PBS+sham (top left), PBS+AP (top right), NT+AP (bottom left) and TNFR1+AP (bottom right) treatment groups. **D).** Percentage of IL-1β immunopositive NP cells; *denotes p<0.05 compared to PBS+sham. **E).** Images of IHC staining for IL-6 expression (brown) in rat NP cells in the PBS+sham (top left), PBS+AP (top right), NT+AP (bottom left) and TNFR1+AP (bottom right) treatment groups. **F).** Percentage of IL-6 immunopositive NP cells; *denotes p<0.05 compared to PBS+sham.

Similarly, IHC staining for IL-1β was present in IVDs from animals in all treatment groups (**Figure 5C**), with more robust IL-1β staining in the PBS+AP and NT+AP treatment groups (**Figure 5C**). Additionally, the percentages of IL-1β immunopositive cells in IVDs from animals in the PBS+AP and NT+AP treatment groups were significantly elevated over the percentages of IL-1β immunopositive cells in the PBS+sham group (**Figure 5D**). Moreover, IVDs from animals in the TNFR1+AP group exhibited significantly reduced percentages of IL-1β immunopositive cells compared to the NT+AP treatment group (**Figure 5D**); however, the percentage of IL-1β immunopositive cells remained significantly elevated over baseline (**Figure 5D**).

IL-6 immunopositive cells were present in IVDs from all treatment groups (**Figure 5E**). The percentage of IL-6 immunopositive cells were significantly elevated over PBS+sham for all experimental treatment groups (**Figure 5F**). Together, these results demonstrate epigenome editing of TNFR1 expression in IVD reduced disc degeneration, recovered disc height, and reduced expression of the inflammatory cytokines TNF-α and IL-1β in an *in vivo* model of disc degeneration and establish CRISPRi epigenome editing of inflammatory cytokine receptor profiles in IVDs as a potential treatment strategy for disc degeneration.

### *In Vivo* CRISPR Based Epigenome Editing of TNFR1 Expression in Rat Caudal IVDs Maintained Disc Height, Prevented Disc Degeneration, and Reduced Inflammation in a TNF-α Injection Model of Disc Degeneration

To further investigate the role of TNF-α signaling through TNFR1 in disc degeneration, we tested the ability of TNFR1 epigenome editing to prevent disc degeneration in an *in vivo* inflammation-driven model of disc degeneration. As above, rat caudal IVDs were injected with either PBS, non-target, or TNFR1-targeting epigenome editing LV vectors (**Figure 6A**). Three weeks after injection, disc degeneration was induced via TNF-α injection. 4 weeks following AP, disc height loss, disc degeneration, and intradiscal inflammatory cytokine levels were evaluated (**Figure 6A**). Again, we confirmed the ability of *in vivo* CRISPRi epigenome editing of the TNFR1 gene promoter to regulate TNFR1 expression during disc degeneration (**Figure 6B**), as the TNFR1 staining intensity of IVDs from animals in the TNFR1-edited plus TNF-α injection (TNFR1+TNF-α) treatment group was significantly decreased when compared to TNFR1 staining intensity of IVDs in the PBS injected plus sham (PBS+sham), and non-target epigenome edited plus TNF-α injected (NT+TNF-α) (**Figure 6B**) treatment groups.

**Figure 6.**
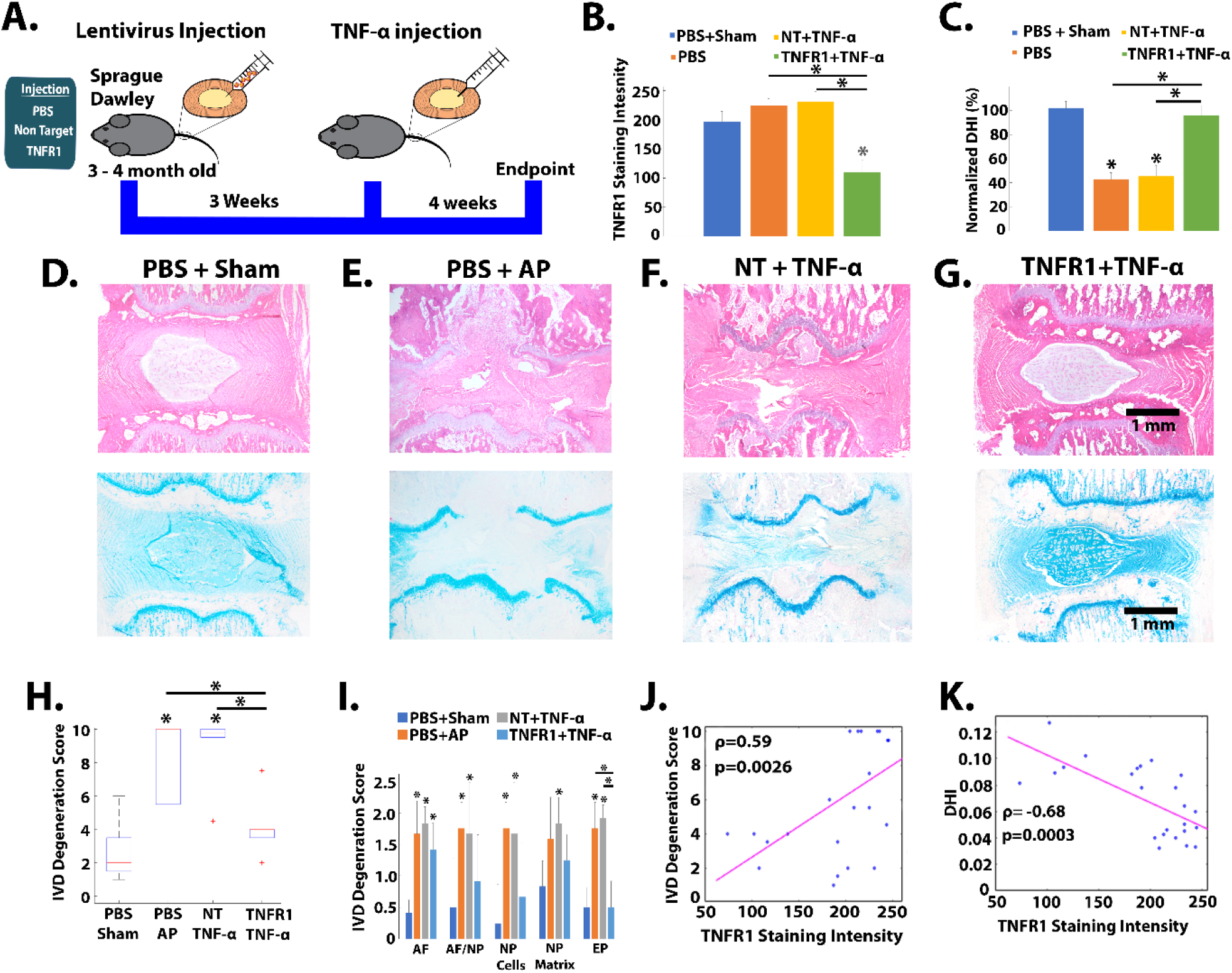
*In vivo* CRISPR-based epigenome editing in rat caudal IVDs prevents disc degeneration and disc height loss in an inflammatory model of disc degeneration. **A).** Schematic of study design: 3-month-old male, Sprague-Dawley rats received caudal intradiscal injection of PBS (n=12 animals), non-target (NT) (n=6 animals) or TNFR1 epigenome editing LV vectors (TNFR1). Three weeks post-injection PBS-injected animals received either sham (n=6) or annular puncture (AP) (n=6), while all NT and TNFR1-injected animals received TNF-α injection. Four weeks following AP, disc height images were obtained, animals were sacrificed, and TNFR1 IHC, histology, and degeneration scoring of IVDs were conducted. **B).** Quantification of TNFR1 immunohistochemistry (IHC) staining intensity of IVDs from animals in the sham (**PBS+sham**), annular puncture (**PBS+AP**), non-target epigenome editing LV injected plus TNF-α injected (**NT+ TNF-α**) and TNFR1 epigenome editing LV vector injected plus TNF-α injected (**TNFR1+ TNF-α)** treatment groups; * denotes p<0.05 compared to PBS+sham. **C).** Normalized disc height index (DHI) calculated from radiographs of caudal IVDs obtained before and after TNF-α injections for animals in the PBS+sham, PBS+AP, NT+TNF-α, and TNFR1+ TNF-α treatment groups; *denotes P<0.05 compared to PBS+sham treatment group. Representative images of H&E (top) and Alcian blue (AB) (bottom) stained histology sections of IVDs from animals in the **D).** PBS+sham, **E).** PBS+AP, **F).** NT+ TNF-α, and **G).** TNFR1+ TNF-α treatment groups. IVD degeneration scores of **H).** total IVDs and **I)**. by IVD region; annulus fibrosus (**AF**), nucleus pulposus (NP), AF and NP border (**AF/NP**), NP cellularity (**NPC**), NP matrix (**NPM**), endplate (**EP**); Scored by blinded observers; * denotes p<0.05 compared to PBS+sham. Spearman correlation modeling of **J).** IVD degeneration score and **K).** DHI as a function of TNFR1 staining intensity. **B**-**I).** n= 6 animals per treatment group.

Next, we investigated the ability of CRISPRi epigenome editing of TNFR1 expression to regulate disc height loss during inflammation driven disc degeneration. The DHI of IVDs from animals in the PBS+AP, and NT+TNF-α (52.4±9%) treatment groups were significantly decreased when compared to the PBS+sham treatment group, indicating significant disc height loss occurred following intradiscal TNF-α injection (**Figure 6C**). However, the DHI of IVDs from animals in the TNFR1+TNF-α (97.3±8.6%, **Figure 6C**) treatment group were maintained at baseline (sham levels). In contrast to annular puncture alone, TNFR1 modulation has shifted TNF-α signaling to a therapeutic response demonstrating the maintenance of all the disc height.

IVD tissue morphology was evaluated from H&E and Alcian blue stained IVD histology sections. IVDs from animals in the PBS+sham group (**Figure 6D**) exhibited healthy IVD tissue architecture characterized by a well-defined NP and an intact NP/AF border, well-defined lamellae structure in the AF, normal disc height, intact end plates, healthy NP cellularity, and robust PG expression in both NP and AF tissue (**Figure 6D**). In contrast, IVDs from animals in the PBS+AP (**Figure 6E**) and NT+TNF-α (**Figure 6F**) treatment groups exhibited classical characteristics of disc degeneration including loss of NP organization and cellularity, loss of AF lamellae structure, endplate fracture, and loss of PG expression; however, IVDs from animals in the TNFR1+TNF-α (**Figure 6G**) exhibited a well-defined NP, with healthy cellularity and distinct NP/AF border, intact endplates, robust PG expression, and recovery of AF lamellae structure. Together, these results demonstrate that TNFR1+TNF-α treatment preserved IVD structure and prevented IVD degeneration under degenerative conditions, and demonstrate the shift of TNA-α to a therapeutic modulator of disc degeneration.

In addition, disc degeneration scores were generated by analysis of IVD histology images by blinded observers. IVDs from animals in the NT+TNF-α treatment group exhibited significantly elevated degeneration scores (8.9±2.2, **Figure 6H**), when compared to IVDs from animals in the PBS+sham group (2.7±1.7). However, disc degeneration scores of IVDs from animals in the TNFR1+TNF-α treatment group (4.2±1.8) were significantly reduced, when compared to IVD degeneration scores from animals in the NT+ TNF-α treatment group and were not significantly different from PBS+sham (baseline) levels (**Figure 6H**).

Analysis of disc degeneration by IVD tissue region revealed IVDs from animals in the PBS+AP or NT+TNF-α treatment groups or both exhibited significantly elevated degeneration scores, compared to IVDs from animals in the PBS+sham treatment group, across all IVD regions (**Figure 6I**); nevertheless, degeneration scores of IVD from animals in the TNFR1+TNF-α group were returned to baseline levels (**Figure 6I**) in the AF/NP, NPC, NPM, and EP regions. Together, these results demonstrate that the combination of TNF-α dosing and TNFR1 knockdown (**Figure 6I**) was most effective at preventing degenerative changes to the NP and endplates of the disc in *in vivo* models of IVD degeneration.

Moreover, Spearman correlation modeling of IVD degeneration scores (**Figure 6J**) and DHI (**Figure 6K**) as a function of TNFR1 expression demonstrated strong correlations between TNFR1 expression, and IVD degeneration or disc height loss, respectively. These results further underscore the relationship between disc degeneration and TNF-α signaling through TNFR1.

In addition to the ability of epigenome editing of TNFR1 expression to ameliorate disc height loss and disc degeneration, we evaluated the ability of CRISPRi based modulation of TNF-α signaling through TNFR1 to regulate inflammatory cytokine expression during disc degeneration. Immunohistochemistry staining of TNF-α in histology sections of IVDs revealed the presence of TNF-α (**Figure 7A**), IL-1β (**Figure 7C**), and IL-6 (**Figure 7E**) immunopositive cells in all treatment groups. The percentage of TNF-α (**Figure 7B**), IL-1β (**Figure 7D**), and IL-6 (**Figure 7F**) immunopositive cells in IVDs from animals in the PBS+AP, and NT+TNF-α treatment groups were significantly elevated over PBS+sham; however, the percentage of TNF-α (**Figure 7B**) and IL-1β (**Figure 7D**) immunopositive cells were significantly reduced compared to the percentage of immunopositive cells in animals from the PBS+AP, and NT+TNF-α treatment groups. The percentage of IL-6 immunopositive cells in IVDs from the TNFR1+TNF-α treatment group were not significantly different from the percentage of IL-6 immunopositive cells in IVDs from animals in the PBS+AP and NT+TNF-α groups (**Figure 7F**). Together, these results demonstrate CRISPRi epigenome editing-based downregulation of TNFR1 expression in the IVD reduced expression of the inflammatory cytokines TNF-α, and IL-1β during disc – degeneration.

**Figure 7.**
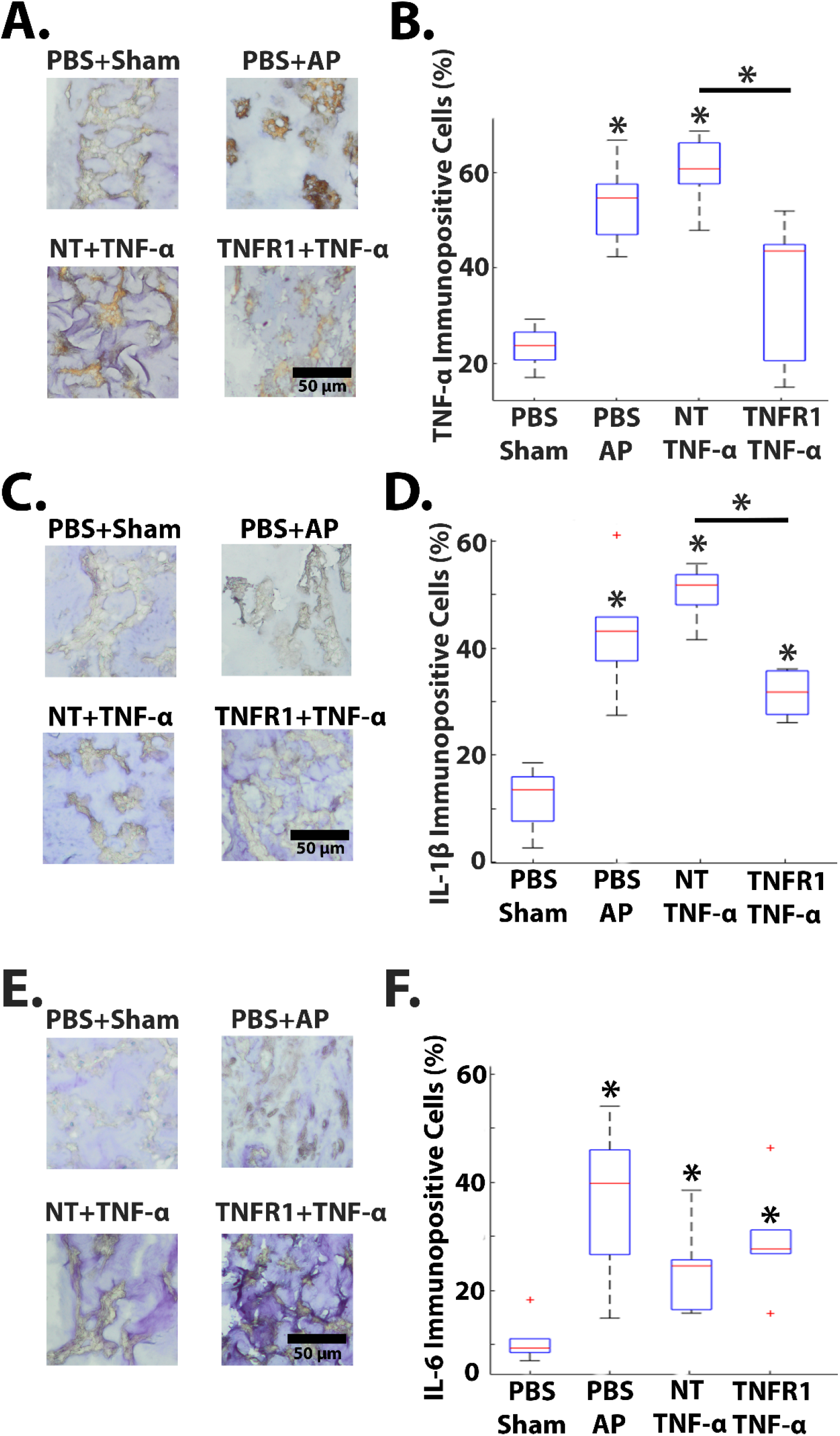
*In vivo* CRISPR epigenome editing of TNFR1 expression in rat caudal IVDs reduced expression of TNF-α, and IL-1β in inflammation model of disc degeneration. **A).** Images of immunohistochemistry (IHC) staining for TNF-α expression(brown) in rat NP cells from IVDs in the PBS injection and sham (**PBS+sham**) (top left), PBS injection and annular puncture (AP) (**PBS+AP**) (top right), non-target epigenome editing LV vector injection and TNF-α (**NT+ TNF-α**) (bottom left), and TNFR1 epigenome editing LV vector injection and TNF-α injection (**TNFR1+ TNF-α**) (bottom right) treatment groups. **B).** Percentage of TNF-α immunopositive NP cells; n=6 animals per group; *denotes p<0.05 compared to PBS+sham **C).** Images of IHC staining for IL-1β expression (brown) in rat NP cells in the PBS+sham (top left), PBS+AP (top right), NT+ TNF-α (bottom left) and TNFR1+ TNF-α (bottom right) treatment groups. **D).** Percentage of IL-1β immunopositive NP cells; n= 6 animals per group; *denotes p<0.05 compared to PBS+sham **E).** Images of IHC staining for IL-6 expression(brown) in rat NP cells in the PBS+sham (top left), PBS+(top right), NT+ TNF-α (bottom left) and TNFR1+ TNF-α (bottom right) treatment groups. **F).** Percentage of IL-6 immunopositive NP cells; n= 6 animals per group; *denotes p<0.05 compared to PBS+sham.

When comparing PBS+Sham, PBS+AP, TNFR1-edited, and TNFR1+TNF-α groups directly (**Figure 8A-D**), a significant increase in blinded degeneration scores was seen in the TNFR1+TNF-α, but not in the TNFR1-edited group when compared to the PBS+AP group (**Figure 8E**). Additionally, a significant decrease shown in normalized DHI (%) between the TNFR1-edited group and the PBS+Sham group, but this was restored in the TNFR1+TNF-α group (**Figure 8F**). Together, these results point to potential therapeutic and degeneration preventative effects in the TNFR1+TNF-α group via TNF-α/TNFR2 signaling.

**Figure 8.**
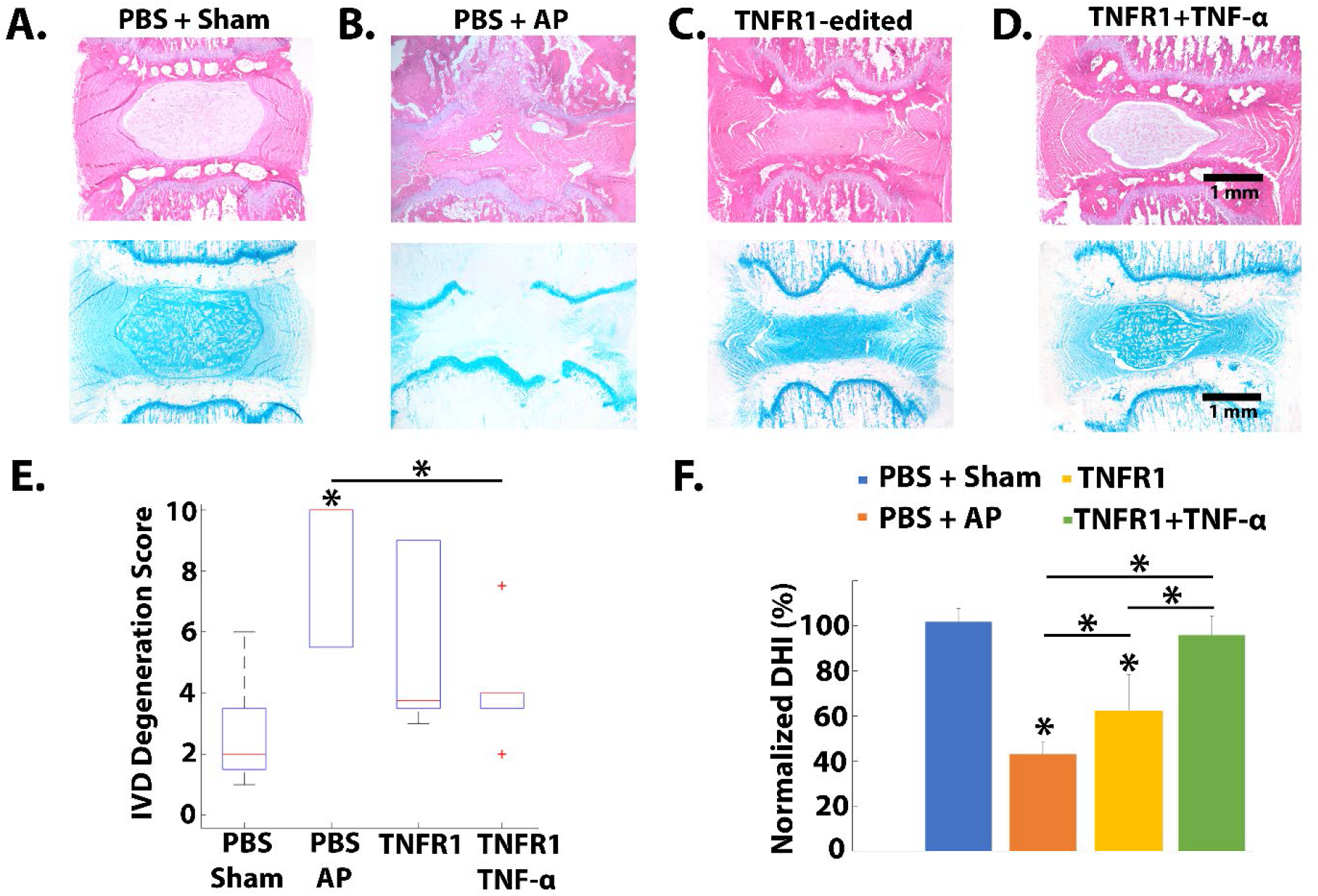
*In vivo* CRISPR-based epigenome editing of TNFR1 and injection of TNF-α in rat caudal IVDs prevents disc degeneration and disc height loss more than editing of TNFR1 alone. Representative images of H&E (top) and Alcian blue (AB) (bottom) stained histology sections of IVDs from animals in the **A).** PBS+sham, **B).** PBS+AP, **C).** TNFR1, and **G).** TNFR1+ TNF-α treatment groups. **E).** IVD degeneration scores of total IVD. **F).** Normalized disc height index (DHI) calculated from radiographs of caudal IVDs obtained before and after TNF-α injections for animals in the PBS+sham, PBS+AP, NT+TNF-α, and TNFR1+ TNF-α treatment groups; *denotes p<0.05 compared to PBS+sham treatment group.

### *In Vivo* CRISPR Based Epigenome Editing of TNFR1 Expression in Rat Lumbar IVDs Reduced Behavioral Pain in a Lumbar IVD Injury Model of Disc Degeneration

To further investigate the therapeutic potential of CRISPR-based TNFR1 epigenome-editing vectors, we explored their ability to reduce painful behavior in an *in vivo* model of lumbar disc degeneration. First, we established our ability to consistently deliver microliter injections to the lumbar spine IVD using a novel, non-surgical, image-guided approach (**Figure 9B**). After optimizing our method, we were able to deliver dye to the center of 9/9 rat cadaver IVDs using our non-surgical method (**Figure 9C**).

**Figure 9.**
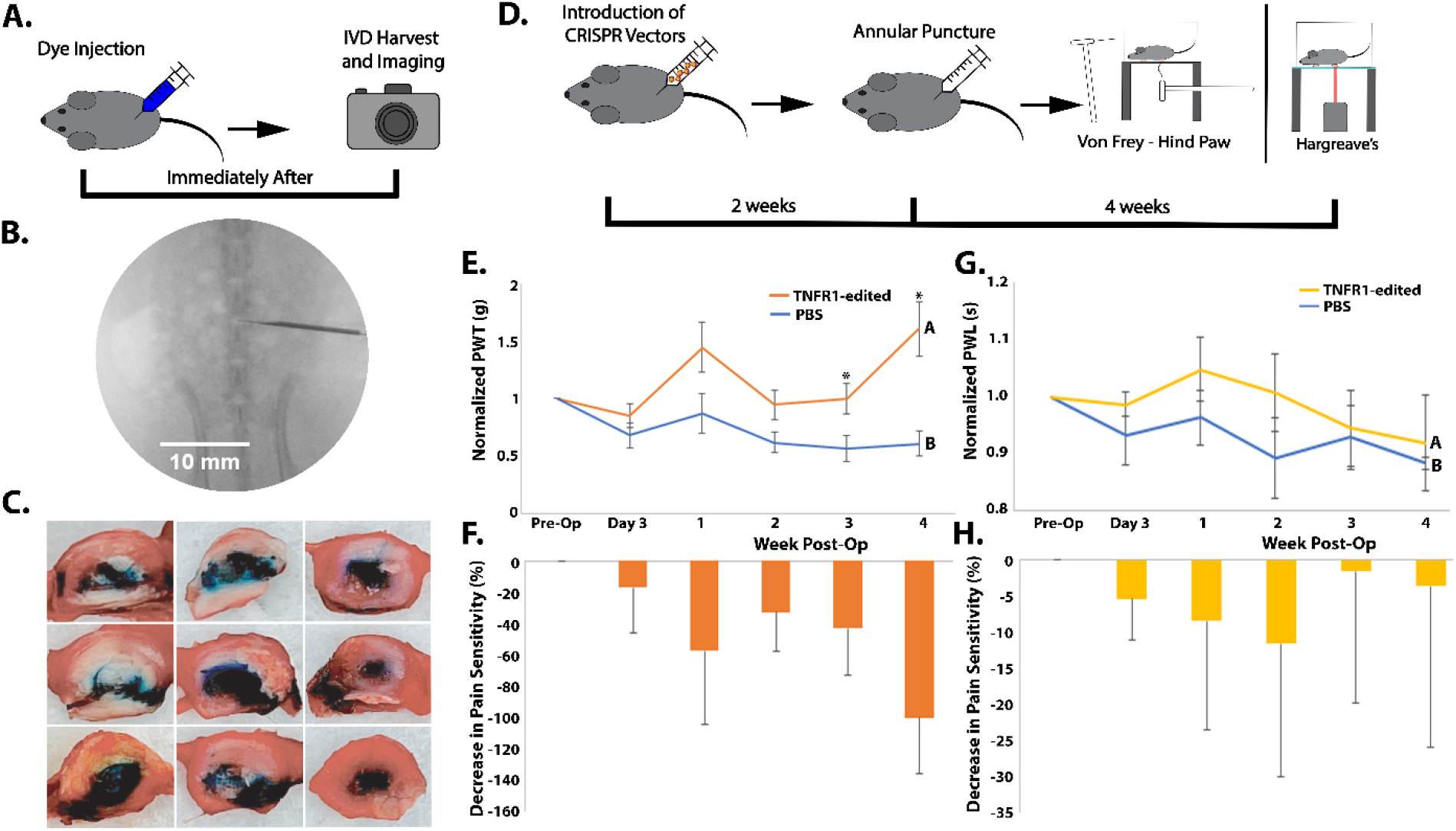
*In vivo* CRISPR epigenome editing of TNFR1 expression in rat lumbar IVDs reduced thermal and mechanical pain sensitivity in annular puncture model of disc degeneration. **A).** Schematic of practice dye injections. Lumbar injections were performed and the discs were immediately harvested for subsequent imaging. **B).** Example c-arm fluoroscope image of lentivirus delivery to IVD using novel approach. **C).** Harvested IVD’s from practice injections showing successful dye injection in nucleus pulposus of IVD. **D).** Schematic of CRISPR epigenome-editing of TNFR1 experiments with behavioral assays: 3-month-old male, Sprague-Dawley rats received lumbar intradiscal injection of PBS (n=7 animals) or TNFR1 epigenome editing LV vectors (TNFR1) (n=7). Two weeks post-injection all animals received annular puncture (AP). Von Frey and Hargreaves behavioral assays were then performed at 3 days post-AP and then weekly for four weeks. **E).** Normalized paw withdrawal threshold (PWT) means of PBS and TNFR1-edited groups using Von Frey behavioral assay. Error bars are standard mean error. **F).** Decrease in mechanical pain sensitivity from Von Frey behavioral assay between PBS and TNFR1-edited groups. Error bars are standard deviation. **G).** Normalized paw withdrawal latency (PWL) means of PBS, TNFR1-, and IL6ST-edited groups using Hargreaves behavioral assay. Error bars are standard mean error. **H).** Decrease in thermal pain sensitivity from Hargreaves behavioral assay between PBS, TNFR1-, and IL6ST-edited groups. Error bars are standard deviation. Groups denoted with different letters indicate statistically significant differences between groups and * denotes statistically significant differences at certain time points, α=0.05.

**Table 1:** Rat weights.

| **Day** | **1** | **7** | **14** | **21** | **28** | **35** | **42** | **49** |
| --- | --- | --- | --- | --- | --- | --- | --- | --- |
| *Caudal Delivery:* |  |  |  |  |  |  |  |  |
| PBS + Sham | 382±15 g | 398±38g | 425±41g | 455±53g | 472±52g | 482±53g | 503±61g | 506±63g |
| PBS + AP | 370±19 g | 390±17g | 435±31g | 467±39g | 487±39g | 507±34g | 508±21g | 552±44 |
| NT + AP | 342±41g | 402±10g | 417±24g | 440±36g | 445±54g | 472±44g | 497±34g | 508±39g |
| NT + TNF-α | 373±31g | 402±28g | 433±40g | 460±56g | 487±52g | 502±52g | 517±63g | 537±74g |
| TNFR1 + AP | 378±26g | 400±18g | 428±26g | 453±22g | 480±40g | 495±47g | 518±53g | 522±46g |
| TNFR1 + TNF-α | 373±25g | 405±23g | 463±27g | 485±26g | 485±32g | 510±43g | 520±32g | 556±64g |
| *Lumbar Delivery:* |  |  |  |  |  |  |  |  |
| PBS + AP | 364±21g | 389±24g | 427±26g | 434±29g | 461±39g | 485±35g | 509±36 | N/A |
| TNFR1 + AP | 369±20g | 397±28g | 436±32g | 443±34g | 472±37g | 500±36g | 517±39g | N/A |
| All Rats | 369±27g | 396±25g | 433±33g | 455±37g | 473±42g | 494±42g | 512±42g | 531±56g |

After practice cadaver injections, we performed our *in vivo* lumbar experiments (**Figure 9D**). Following acclimation sessions, pre-procedural baseline data was collected for each assay on each animal. After baseline data collection, lumbar IVDs were injected with either PBS or TNFR1-targeting epigenome-editing LV vectors. Two weeks after injection, disc degeneration was induced via AP. Von Frey and Hargreaves behavioral assays were then performed at 3 days and 1-week post-AP and then weekly for 4 weeks post-AP.

Our Von Frey showed a significant reduction in normalized paw withdrawal threshold (PWT) between PBS and TNFR1-targeting groups and at time points at 3- and 4-weeks post-AP (**Figure 9E**). These results show a reduction in induced mechanical pain sensitivity in our degenerative lumbar IVD model (**Figure 9F**). In addition, our Hargreaves behavioral data showed a significant reduction in normalized paw withdrawal latency (PWL) of the TNFR1-targeting group compared to the PBS group (**Figure 9G**). These results demonstrate a change in thermal pain sensitivity in our lumbar injury model (**Figure 9H**). Together, these results demonstrate CRISPRi epigenome editing-based downregulation of TNFR1 expression in the IVD reduces pain sensitivity from disc degeneration.

## DISCUSSION

Despite the prevalence of disc degeneration and its associated pathologies^2, 4, 59^, and the enormous socio-economic cost generated by their treatment^5, 60^, effective treatment strategies capable of slowing the progression of disc degeneration remain elusive. Currently, clinical treatments focus on palliative care, and surgical interventions remain sub-optimal for symptom relief and inadequate at preventing further disc degeneration. Recently, the advent of CRISPR^61–64^ epigenome-editing technology has provided new tools to allow the development of therapeutics that specifically target the molecular signaling pathways causing disc degeneration.

Pathologically high levels of TNF-α are prevalent in degenerative discs^19, 28, 29, 32, 34^ and TNF-α signaling has been implicated as an underlying mechanism that drives disc degeneration^31–35^. Previously, we have demonstrated the ability of CRISPRi epigenome editing of TNFR1 to mediate inflammatory cytokine-driven pathological changes in *in vitro* models of disc degeneration^51, 56^. In this study, we investigated the therapeutic potential of LV-delivered CRISPRi epigenome-editing vectors targeting the TNFR1 gene promoter to slow disc degeneration in rodent *in vivo* models of disc degeneration.

In order to be a viable clinical treatment strategy, CRISPRi epigenome editing vectors must be delivered to the IVD effectively, be expressed locally in the IVD, and maintain long-term downregulation of target gene expression. In this study, *ex vivo* bioluminescent imaging of rat caudal IVDs injected with luciferase-expressing LV vectors demonstrated sequestration of LV expression within the target IVD (**Figure 1B**) and no expression in vital organs or adjacent tissues (**Figure 1C**). Combined with previous studies demonstrating the avascular nature of the IVD^65^, and LV expression localized to the injection site ^66, 67^, these results suggested expression of CRISPRi epigenome editing LV, and hence, gene regulation will be localized within the injected IVD.

Once we established the ability to safely deliver LV vectors to the rat caudal IVDs, we investigated the ability of CRISPRi LV vectors to regulate TNFR1 expression *in vivo*. Previously, we demonstrated the ability of CRISPRi based therapeutics to downregulate TNFR1 expression *in vitro*^51, 52, 56^, maintain TNFR1 downregulation of TNFR1 expression under inflammatory conditions^51, 52^, and modulate TNF-α/TNFR1 pro-inflammatory signaling-driven cellular processes^51, 52, 56^. In this study, we delivered TNFR1-targeted CRISPRi-based therapeutics to rat caudal IVDs and measured their ability to regulate TNFR1, at time points of 3 weeks and 3 months after delivery, via IHC. Downregulation of TNFR1 expression was demonstrated 3 weeks after therapeutic delivery (**Figure 2**) and maintained for 3 months post-treatment (**Figure 3**), while healthy IVD tissue morphology was preserved. These results demonstrated the ability of CRISPRi-based therapeutic treatments to maintain long-term downregulation of TNFR1 expression *in vivo*, ergo illustrating delivery of CRISPRi-based therapeutics to the IVD as a potential treatment strategy for disc degeneration by regulating inflammatory signaling cascades in the degenerative IVD.

Previous studies have demonstrated the role of TNF-α in the progression of disc degeneration^26, 55, 68^, and that treatments blocking TNF-α can mediate disc degeneration^68, 69^; however, these treatments with these agents have short term effects and would require multiple disc punctures^41, 69^. In this study, we demonstrated the ability of our CRISPRi-based therapeutics to provide long-term downregulation of TNFR1 expression and slow disc degeneration following annular puncture. Utilizing this strategy, we hypothesized we could shift TNF-α signaling from the pro-inflammatory TNFR1-mediated pathway^35^ to anti-inflammatory and immunomodulatory TNFR2-mediated signaling pathways^35, 40^, simultaneously blocking inflammatory signaling and encouraging anti-inflammatory signaling to attenuate disc degeneration. To test this hypothesis, we tested the ability of TNFR1-targeting CRISPRi therapeutics to reduce disc degeneration following a TNF-α injection (**Figure 6A**). IVDs that received TNF-α injection following treatment with TNFR1-targeted CRISPRi-based therapeutics exhibited no loss of disc height (**Figure 6C**), maintained healthy IVD tissue architecture (**Figure 6G**) and disc degeneration scores (**Figure 6H**), and reduced inflammatory cytokine expression to a greater degree than IVDs that received TNFR1-targeted CRISPRi therapeutic treatment alone. Together, these results demonstrate the ability of IVD treatment with TNFR1-targeting CRISPRi-based therapeutics to prevent disc degeneration and modulation of inflammatory receptors can cause significant beneficial shifts in inflammatory cytokine effects *in vivo*.

After establishing the therapeutic potential of our CRISPRi-based epigenome-editing in a caudal model of disc degeneration, we sought to move delivery to the lumbar spine and explore the effects that these vectors have on behavioral pain. Despite the difficulty of accessing the lumbar spine IVDs, we were able to utilize the same novel non-surgical, image-guided procedures used in the caudal spine for both the LV delivery and the AP in the lumbar spine (**Figure 9B**). We demonstrated the ability to consistently deliver to the lumbar spine IVD on rat cadavers using this method in 9/9 discs in our final practice procedure (**Figure 9C**).

In our *in vivo* lumbar delivery studies, we demonstrated the ability to significantly reduce behavioral pain with our TNFR1-targeted CRISPRi therapeutics. Our behavioral data show a significant reduction in thermal and mechanical pain sensitization over 4 weeks post-annular puncture between TNFR1-targeted delivery and PBS delivery groups (**Figure 9E-H**). These behavioral data establish the capability of our therapeutics to reduce induced pain sensitivity in addition to modifying degenerative disc disease progression. The observed pain behavior modifications may be due to the observed maintenance of IVD structure (**Figures 3G, 4G, 6G)**, the overall reduction of inflammatory cytokine presence (**Figures 5, 7)**, or TNFR1 modification of innervating neurons in the disc (**Figures 2C, 3C)**. All of these are consistent with our previous in vitro model work with human cells and tissue demonstrating the reduction of inflammatory-driven cell apoptosis/degeneration^57^ and neuron sensitization^58, 70^.

Overall, we have demonstrated the safe *in vivo* delivery of CRISPRi therapeutics to the IVD and their ability to maintain long-term inflammatory receptor modulation. In addition, we demonstrate IVD treatment with TNFR1-targeting CRISPRi therapeutics preserved acute and chronic changes to IVD height, disc degeneration progression, inflammation, and pain sensitization in multiple *in vivo* rodent models of disc degeneration. These results elucidate TNFR1 as a potential target for therapeutics designed to slow the progression of disc degeneration and establish CRISPRi-based epigenome editing of TNFR1 expression in the IVD as a potential treatment strategy for disc degeneration. Additionally, this work demonstrates that inflammatory receptor targeting can protect beneficial inflammatory signaling that direct cytokine targeting eliminates. Future work will investigate the efficacy of CRISPRi-based therapeutic treatment of the IVD for broader regulation of inflammatory pathways in the IVD and for treating low back pain in animal models of disc degeneration.

## METHODS

### Experimental Overview

In this study, a set of four *in vivo* experiments were conducted to investigate the ability of intradiscal delivery of CRISPR-based epigenome editing lentiviral vectors to regulate disc degeneration and pain in *in vivo* rodent models of inflammatory disc degeneration. In the first set of experiments, rat caudal IVDs received intradiscal injection of lentiviral vectors containing luciferase, non-target, or TNFR1-targeting epigenome-editing vectors. Bioluminescent imaging was conducted to determine the ability to deliver and sequester lentiviral vectors in the target IVD. In addition, immunohistochemistry staining and morphology-based disc degeneration scoring was performed to investigate the ability of TNFR1 epigenome editing of the IVD to downregulate TNFR1 expression, while maintaining healthy disc morphology. A subsequent set of experiments were conducted to investigate the potential of CRISPR epigenome editing of the IVD to produce long-term downregulation of TNFR1 expression in the IVD without altering healthy disc morphology. A third set of experiments was performed to evaluate the ability of CRISPR epigenome editing of TNFR1 expression to regulate disc degeneration, disc height loss, and inflammation in an *in vivo* rodent model of disc degeneration. A final set of experiments was performed the investigate the ability of CRISPR epigenome editing of TNFR1 expression to regulate thermal and mechanical pain sensitization in an *in vivo* rodent model of lumbar spine disc degeneration.

### Animals

All procedures were performed with approval of the University of Utah Institutional Animal Care and Use Committee (IACUC). A total of 108 skeletally mature male (10-12 week) Sprague Dawley rats were used in this study. Animals were allowed free movement in their home cages and were housed two per cage throughout the study. Rats were allowed *ad libitum* access to food and water. To ensure animal health, rats underwent routine health and wellness exams and were weighed on a weekly basis. Rats were maintained at 72-74°F and 12/12 hour light/dark cycle (6am-6pm light on).

### Intradiscal Injection of Lentiviral Vectors

Rat IVD intradiscal injections were performed under sterile conditions and general anesthesia (2% isoflurane inhalation). IVDs were visualized via radiographic imaging and IVD levels were identified via anatomical landmarks. For the caudal IVD studies, a 31-gauge needle was positioned in the center of the disc space using fluoroscopic guidance (**Figure 1A**) and intradiscal injection of PBS, luciferase-expressing lentiviral vector, and non-target or TNFR1-targeting CRISPR epigenome editing lentiviral vectors (5μL) was conducted. For the lumbar IVD study, a 30-gauge needle fed through a 20-gauge cannula was used for LV delivery was used to aid in delivery. Following injection, rats were closely monitored for complications and allowed *ad libitum* access to food and water.

### IVD Puncture and TNF-α Injection Injury

Rat IVD puncture and TNF-α injection injuries were performed under sterile conditions and general anesthesia (2% isoflurane inhalation). IVDs were visualized using radiographic imaging and IVD levels were identified via anatomical landmarks. In addition, radiographs obtained during **Intradiscal Injection of Lentiviral Vectors** experiments were referenced to ensure IVD injury of disc receiving previous injection. For the caudal IVD studies, a 26-gauge needle was positioned in the center of the disc space, and intradiscal injection of PBS or TNF-α (0.25ng in 2.5μL PBS) was performed^69, 71^. For the lumbar spine studies, a 25-gauge needle was used for AP. Following IVD injuries, rats were closely monitored for complications and allowed *ad libitum* access to food and water.

### *Ex Vivo* Lentiviral Vector Expression Bioluminescent Imaging

Ex vivo imaging was performed, following a previously published method^50^ to determine expression of intradiscally injected lentiviral vectors. At sacrifice, a tail segment containing the injected caudal IVD, a control tail segment, heart, lungs, liver, spleen, and kidneys were harvested for immediate luminescent imaging. Following harvest, tissues were placed in 12-well plates containing 300μg/ml D-luciferin in Dulbecco’s PBS and incubated for 5 minutes. After incubation, tissues were imaged on an IVIS imaging system. ROI containing each tissue was measured as photons/second/centimeter squared.

### Radiographic Disc Height Measurement

IVD heights of non-injured and injured IVDs were measured from radiographs obtained while animals were under general anesthesia. Pre-injury radiographs were obtained immediately before disc puncture and post-injection radiographs were obtained on the day preceding animal sacrifice. Disc height index (DHI) was calculated and normalized to pre-injury DHI using a previously reported method^72^.

### Tissue Harvesting

At the end of experiments, rats were sacrificed via carbon dioxide inhalation. Immediately following sacrifice, rat caudal spines were harvested, fixed, decalcified, embedded in formalin, and sectioned (5μm).

### Disc Degeneration Grading

IVD sections were stained with Alcian blue or hematoxylin/eosin (H&E) to visualize IVD morphology and bright-field microscopy imaging was conducted. IVD degeneration was assessed using a previously reported semi-quantitative degeneration grading scale^73^. The combined degeneration scale (0-10) was comprised of the five following categories: 1) annulus fibrous, 2) border between the AF and NP, 3) NP cellularity, 4) NP matrix, and 5) endplate. Each category received a score between 0 and 2, with a score of 0 indicating a normal (healthy) morphology and a score of 2 indicating severe degeneration. All IVD sections were scored by two observers blinded to the experimental groups.

### Measurement of Intradiscal TNFR1, TNF-α, IL-1β, and IL-6 Expression

Intradiscal expression of the inflammatory cytokine receptor TNFR1 and the pro-inflammatory cytokines TNF-α, IL-1β, and IL-6 was determined using immunohistochemistry. IVD sections from each rat were deparaffinized, rehydrated, blocked, and incubated overnight with either rabbit polyclonal primary antibody against TNFR1, rat TNF-α, rat IL-1β, rat IL-6, or isotype control antibody. Following overnight incubation, sections were incubated with horseradish peroxidase-conjugated anti-rabbit secondary antibody for 30 minutes, followed by a 10-minute incubation in 3% hydrogen peroxide. To visualize immunoreactivity, sections were treated with a horseradish-peroxidase substrate. Subsequently, sections were counterstained with toluidine blue, dehydrated, and evaluated using bright-field microscopy.

### Von Frey and Hargeaves Behavioral Assays

Fourteen skeletally mature male rats were acclimated to the experimenter and the Von Frey mesh bottom (Stoelting) and Hargreaves glass bottom (IITC Life Science) testing apparatuses for three sessions before beginning data collection. For the Von Frey assay, the up-down method and tables were used^74, 75^ with a set of nine filaments (0.4 g, 0.6 g, 1 g, 2 g, 4 g, 6 g, 10 g, 15 g). Five consecutive non-responses on a particular paw was assigned the value of 17.9 g. Positive or negative responses were recorded for each paw separately until four responses were recorded after the first change in response, for a maximum of 9 stimulations per paw. A minimum of 2 minutes was given between each stimulation on the same paw. For Hargreaves, active intensity of the heat source was set to 35% with an automatic cutoff time of 20 seconds. Three recordings were done on each paw, with a minimum of 5 minutes between stimulations on the same paw. For each data collection session, animals were allowed to acclimate in the behavioral apparatuses until self-grooming behavior had largely stopped.

Following acclimation sessions, baseline data was collected for each assay on each animal. After baseline data collection, under anesthesia, lumbar IVDs were injected with 5 µL of either PBS (n=7) or TNFR1-targeting epigenome-editing LV vectors (n=7) (**Figure 9D**) as described in **Intradiscal Injection of Lentiviral Vectors**. Two weeks after injection, disc degeneration was induced via AP as described. Von Frey and Hargreaves behavioral assays were then performed at 3 days and 1-week post-AP and then weekly for 4 weeks post-AP (**Figure 9E-H**). Experimenter was blinded for entirety of behavioral data collection.

### Statistical Analysis

Luminescence, IVD degeneration scoring, intradiscal IHC intensity, disc height index (DHI), and percentage intradiscal immunopositive cells data were all analyzed using one-way ANOVA with Tukey’s test as post hoc comparison. Behavioral data was analyzed using a two-way ANOVA with Tukey’s post hoc test. The correlation between DHI, IVD degeneration score, percentage intradiscal TNF-α, IL-1β, and IL-6 immunopositive cells, and intradiscal TNFR1 expression were analyzed using Pearson’s correlation.

## Acknowledgements

Research reported in this publication was supported by National Institute of Arthritis and Musculoskeletal and Skin Diseases (NIAMS) of the National Institutes of Health under award number R01AR074998.

